# A curated knowledgebase enabling a network perspective on endocrine disrupting chemicals and their biological systems-level perturbations

**DOI:** 10.1101/619163

**Authors:** Bagavathy Shanmugam Karthikeyan, Janani Ravichandran, Karthikeyan Mohanraj, R.P. Vivek-Ananth, Areejit Samal

## Abstract

Human well-being can be affected by exposure to several chemicals in the environment. One such group is endocrine disrupting chemicals (EDCs) that can perturb the hormonal homeostasis leading to adverse health effects. In this work, we have developed a detailed workflow to identify EDCs with supporting evidence of endocrine disruption in published experiments in humans or rodents. Thereafter, this workflow was used to manually evaluate more than 16000 published research articles and identify 686 potential EDCs with published evidence in humans or rodents. Importantly, we have compiled the observed adverse effects or endocrine-specific perturbations along with the dosage information for the potential EDCs from their supporting published experiments. Subsequently, the potential EDCs were classified based on the type of supporting evidence, their environmental source and their chemical properties. Additional compiled information for potential EDCs include their chemical structure, physicochemical properties, predicted ADMET properties and target genes. In order to enable future research based on this compiled information on potential EDCs, we have built an online knowledgebase, **D**atabase of **E**ndocrine **D**isr**u**pting **C**hemicals and their **T**oxicity profiles (DEDuCT), accessible at: https://cb.imsc.res.in/deduct/. After building this comprehensive resource, we employed a network biology approach to study the chemical space of EDCs and its potential link to the biological space of target genes of EDCs. Specifically, we have constructed two networks of EDCs using our resource based on similarity of chemical structures or target genes. Ensuing analysis of these two networks revealed that EDCs can differ both in their chemical structure and set of target genes. Though our detailed results highlight potential challenges in developing predictive models for EDCs, the compiled information in our resource will undoubtedly enable future research in the field, especially, those focussed towards mechanistic understanding of the systems-level perturbations caused by EDCs.

## 1. Introduction

In the last century, industrial advances has led to the rapid synthesis and commercialization of more than 80000 chemicals, however, only a small fraction of these chemicals have been tested for their safety or toxicity concern (Futran Fuhrman et al., 2015; Schug et al., 2013). Humans are exposed to these chemicals in their daily life in the form of consumer products including personal care products, pharmaceuticals, food additives, pesticides and insecticides (Meeker, 2010; Mezcua et al., 2012; Muncke, 2011; WHO/UNEP, 2013). In recent times, certain groups of these chemicals have received serious attention from scientists, regulatory agencies and public due to their potential safety concern. Endocrine disrupting chemicals (EDCs) is one such group listed under the chemicals of emerging concern (Futran Fuhrman et al., 2015). EDCs interfere with the normal functioning of the human endocrine system and can lead to adverse effects related to reproduction, development, metabolism, immune system, neurological system, liver or hormone-related cancers (Solecki et al., 2017; Swedenborg et al., 2009; WHO/UNEP, 2013). EDC exposure can alter hormonal imbalance in humans through different mechanisms. For example, EDCs can mimic the natural hormones and bind to their respective nuclear receptors either as an agonist or an antagonist (Schug et al., 2013; Zoeller et al., 2012). So far there is a lack of biological systems or pathway level understanding of the different mechanisms via which specific EDCs alter the hormonal homeostasis.

For the risk assessment of EDCs, an important limitation is the lack of availability of validated test systems for their identification (Solecki et al., 2017; Zoeller et al., 2012). This has hampered both researchers and policymakers to reach a consensus agreement on identification of EDCs and the characterization of their endocrine disruption mechanisms (Solecki et al., 2017; Zoeller et al., 2012). In this direction, Solecki et al. (Solecki et al., 2017) have outlined a detailed consensus statement on the scientific principles that can form a basis for the identification of EDCs and their disruption mechanism. Furthermore, the scientific statements by the endocrine society (Diamanti-Kandarakis et al., 2009; Gore et al., 2015; Zoeller et al., 2012) provide endocrine principles for better understanding of disruption mechanisms by EDCs.

Given the potential risk from EDCs in our environment, there have been multiple efforts towards their compilation which include the World Health Organization (WHO) report (WHO/UNEP, 2013), The Endocrine Disruption Exchange (TEDX; https://endocrinedisruption.org/), EDCs Databank (Montes-Grajales and Olivero-Verbel, 2015) and Endocrine Disruptor Knowledge Base (EDKB) (Ding et al., 2010). However, these existing resources on potential EDCs consider evidence for endocrine disruption upon exposure from disparate types of published studies including *in vivo*, *in vitro*, *in silico*, environmental monitoring and epidemiological studies. Moreover, these existing resources on potential EDCs do not systematically compile the observed adverse effects specific to endocrine disruption in supporting published experiments.

In this direction, we have developed a detailed workflow (Figure 1) to identify potential EDCs from published research articles containing supporting experimental evidence for endocrine-specific perturbations in humans or rodents. Using this workflow, we processed more than 16000 published research articles to manually compile 686 potential EDCs with supporting evidence of endocrine-specific perturbations from published experiments in humans or rodents. Importantly, the compiled list of observed adverse effects or endocrine-specific perturbations in supporting published experiments for the 686 EDCs were manually curated, unified and standardized into a list of 514 endocrine-mediated endpoints spanning 7 systems-level perturbations (Methods; Figure 2). In contrast to existing resources, another unique feature of our work is the compilation and standardization of the dosage information at which endocrine-mediated effects were observed upon individual EDC exposure in published experiments (Methods). Moreover, the 686 EDCs were classified based on the type of supporting evidence in published experiments, their environmental source and their chemical classification (Methods; Figures 3 and 4). Lastly, we have also compiled additional detailed information for each EDC such as its two-dimensional (2D) and three-dimensional (3D) structure, physicochemical properties, molecular descriptors, predicted ADMET properties and experimentally inferred target genes. In order to widely share the compiled information on 686 potential EDCs and enable basic research towards elucidation of systems-level perturbations caused by them, we have also created a webserver, **D**atabase of **E**ndocrine **D**isr**u**pting **C**hemicals and their **T**oxicity profiles (DEDuCT), which is accessible at: https://cb.imsc.res.in/deduct/.

**Figure 1:**
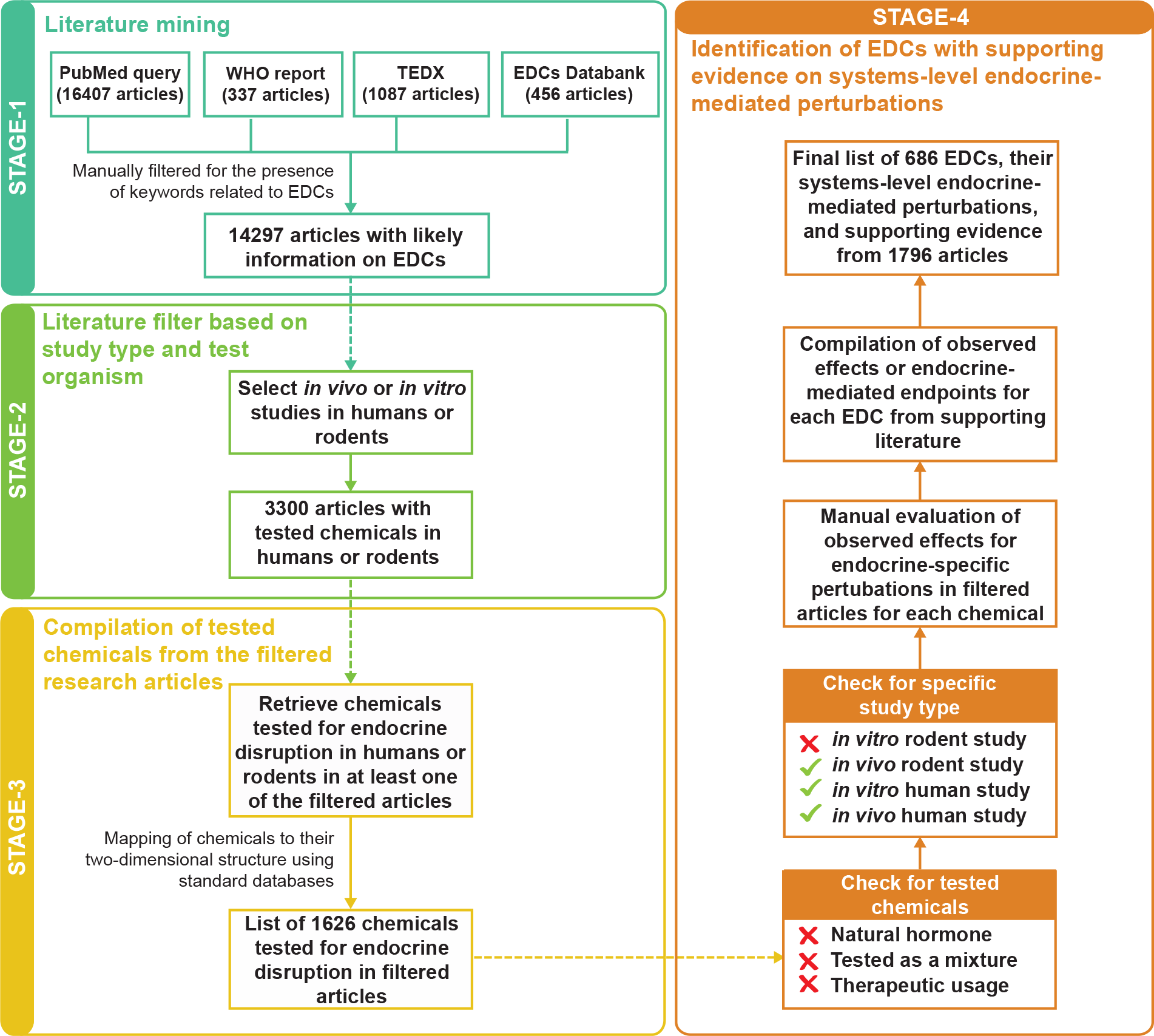
Detailed workflow with four stages to identify potential EDCs from published research articles containing supporting experimental evidence of systems-level endocrine-mediated perturbations in humans or rodents.

**Figure 2:**
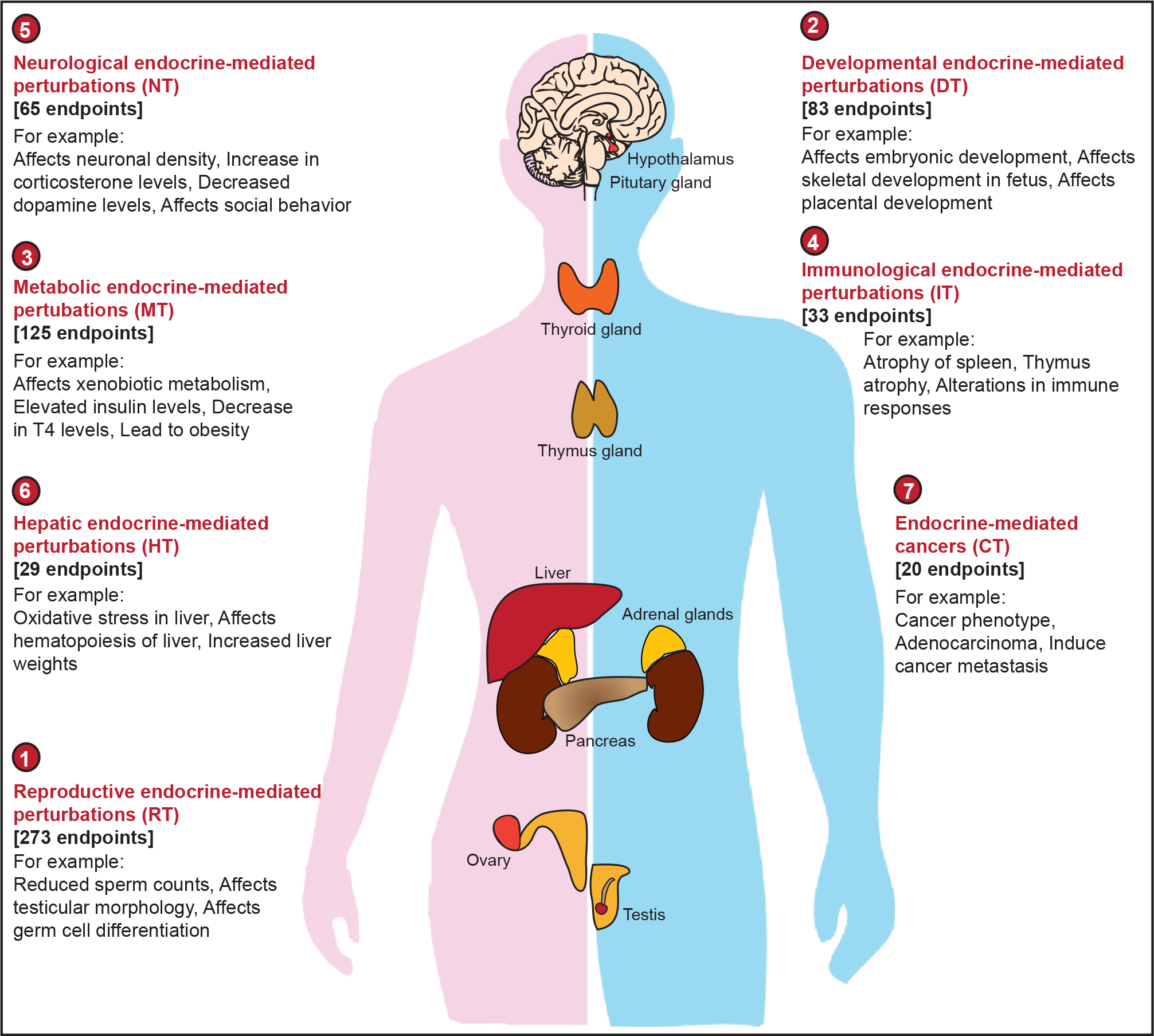
Schematic figure depicting the classification of the 514 endocrine-mediated endpoints into 7 systems-level perturbations. Note that this classification of endpoints into systems-level perturbations is overlapping, that is, a given endpoint may fall into more than one systems-level perturbations.

**Figure 3:**
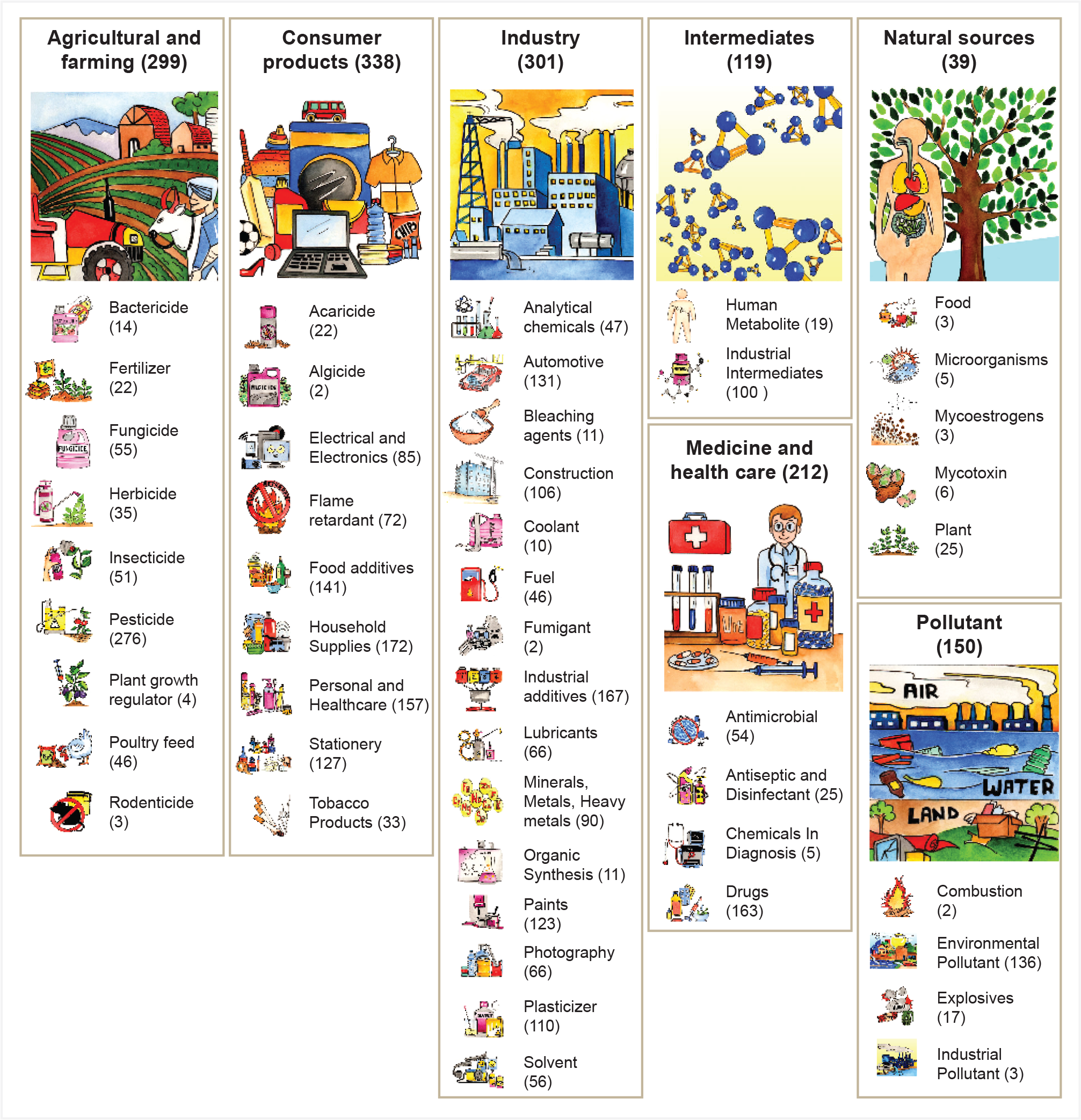
Classification of the 686 EDCs into 7 broad categories and 48 sub-categories based on their source in the environment. In this figure, the number of EDCs in each category or sub-category is reported within the parenthesis. Note that this environmental source-based classification is overlapping, that is, a given EDC may belong to multiple categories or sub-categories.

**Figure 4:**
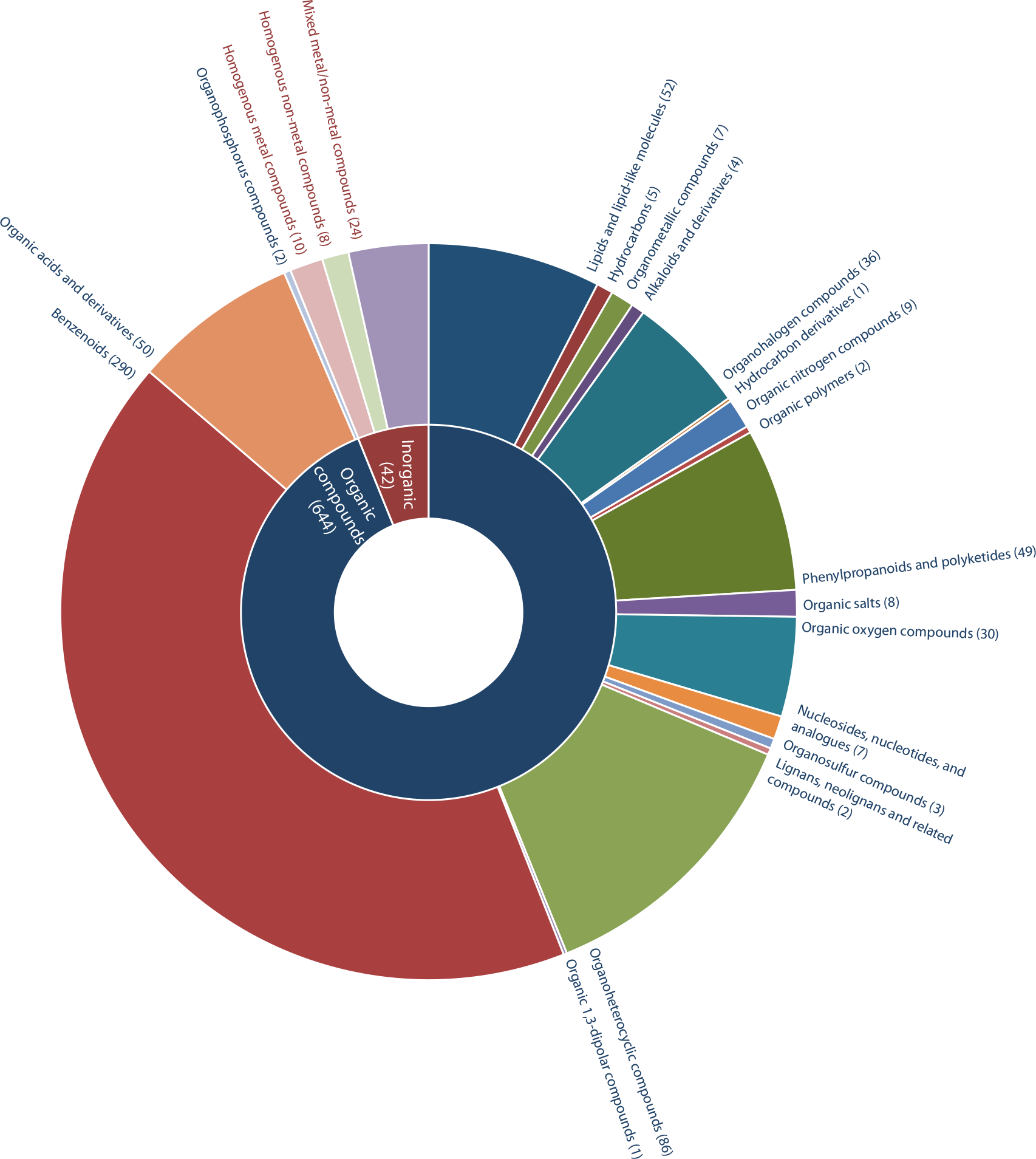
Classification of the 686 EDCs into chemical kingdoms and chemical super-classes using ClassyFire. Of the 686 EDCs, 644 are organic and 42 are inorganic compounds. The 644 organic EDCs can be further classified into 19 super-classes while the 42 inorganic EDCs fall into 3 super-classes. The number of EDCs in each super-class is reported within the parenthesis.

An important motivation for creating this large-scale resource on potential EDCs was to enable network biology approaches (Barabasi et al., 2011; Barabasi and Oltvai, 2004; Zhou et al., 2014) towards a better understanding of the link between the underlying chemical space of EDCs and biological space of target genes or perturbed pathways (Dobson, 2004; Lipinski and Hopkins, 2004). In this direction, we have used our resource to construct two different networks for EDCs, namely, the chemical similarity network and the target similarity network (Methods; Figure 5). Subsequent analysis of these networks revealed the diversity of the chemical space of EDCs and its associated space of target genes. Our results expose future challenges in developing predictive computational systems toxicology models (Kavlock et al., 2008; Merlot, 2010) for EDCs. Altogether, DEDuCT is a large-scale resource on 686 potential EDCs with supporting evidence of endocrine-mediated perturbations and dosage information from published experiments in humans or rodents, and the compiled information will contribute to the future research in the field of computational systems toxicology.

**Figure 5:**
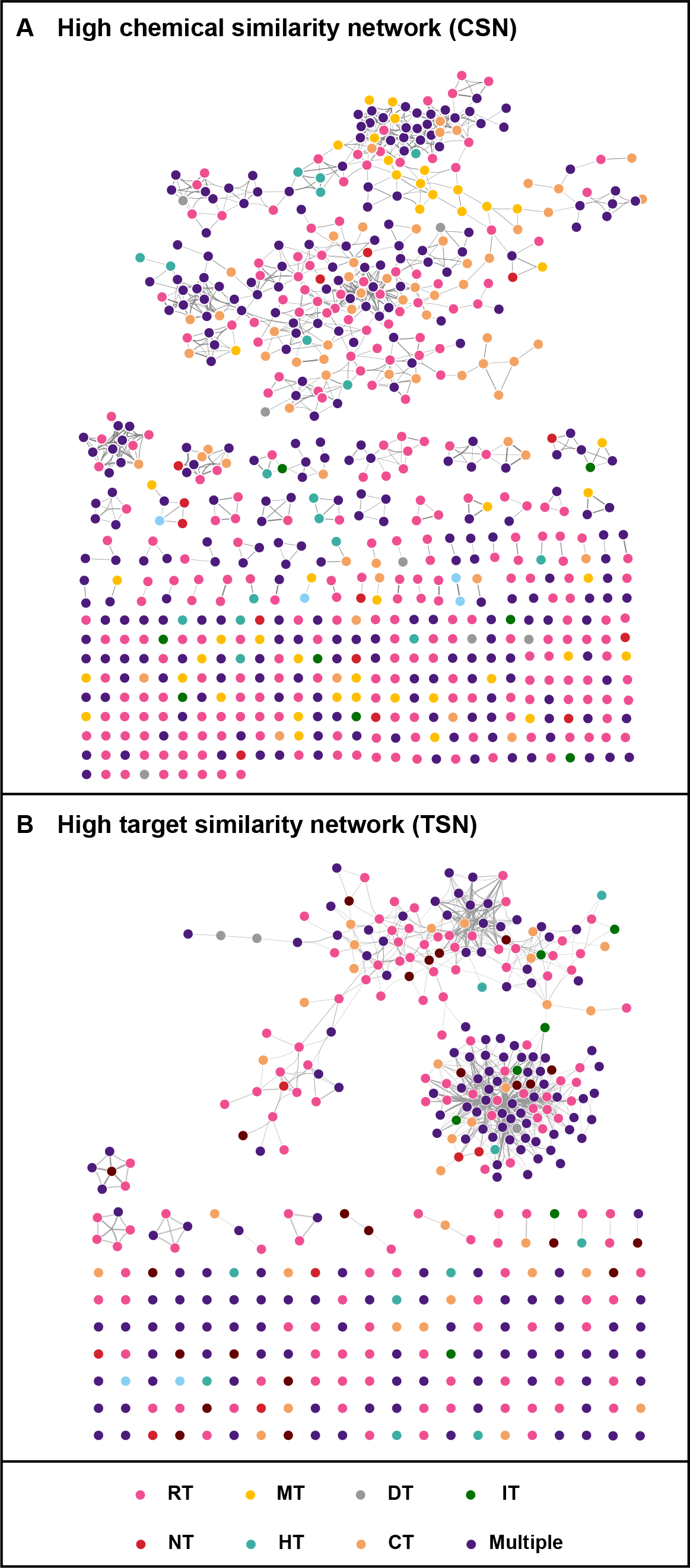
Network of EDCs based on chemical similarity and gene target similarity. **(A)** Network visualization of high chemical similarity network (CSN) of 686 EDCs. **(B)** Network visualization of high target similarity network (TSN) of 383 EDCs. The high TSN was constructed for 383 EDCs which have information on their target genes from ToxCast assays. The legend at the bottom of this figure gives the colour code for nodes or EDCs in CSN and TSN which is based on the 7 systems-level perturbations, namely, Reproductive (RT), Developmental (DT), Metabolic (MT), Immunological (IT), Neurological (NT), Hepatic (HT) and Endocrine-mediated cancer (CT), associated with EDCs in DEDuCT. Note that if an EDC is associated with more than one systems-level perturbations then its colour is given by Multiple. Moreover, the thickness of the edges in CSN and TSN are based on their edge weights given by Tanimoto coefficient and Jaccard index, respectively (Methods).

## 2. Methods

### 2.1 Workflow for the identification of EDCs

Based on the consensus statement by Solecki et al. (Solecki et al., 2017) and the scientific statement by the endocrine society (Diamanti-Kandarakis et al., 2009; Gore et al., 2015; Zoeller et al., 2012), we have developed a detailed flowchart to identify EDCs from published research articles containing supporting experimental evidence of systems-level endocrine-mediated perturbations in humans or rodents (Figure 1). Our workflow for the identification of EDCs can be further subdivided into four stages which are described below.

#### 2.1.1 Literature mining

In stage 1, we performed an extensive literature search to compile 14297 published research articles which are likely to contain information on EDCs (Figure 1).

Firstly, we mined PubMed (https://www.ncbi.nlm.nih.gov/pubmed/) using the following keyword search:

> “EDCs” OR “EDC” OR (“endocrine” AND “disrupt”) OR (“disrupt” AND “endocrine”) OR “endocrine disruptors” OR “endocrine-disruptors” OR “endocrine disruptor” OR “endocrine-disruptor” OR “endocrine disrupters” OR “endocrine-disrupters” OR “endocrine disruption” OR “endocrine-disruption” OR “endocrine disruptive” OR “endocrine-disruptive” OR “endocrine disrupting” OR “endocrine-disrupting” OR “endocrine disrupter”

The above query was designed to filter abstracts on EDCs from PubMed, and this keyword search in February 2018 led to 16407 research articles. Secondly, we compiled research articles from three existing resources on EDCs, namely, the WHO report (WHO/UNEP, 2013), TEDX (https://endocrinedisruption.org/) and EDCs Databank (Montes-Grajales and Olivero-Verbel, 2015) (http://edcs.unicartagena.edu.co/). Specifically, the WHO report, TEDX and EDCs Databank captured information from 337, 1087 and 456 research articles, respectively. Subsequently, we manually filtered the compiled abstracts from PubMed query, WHO report, TEDX and EDCs Databank for the presence of keywords related to EDCs, and this filtration led to 14297 research articles at the end the stage 1 (Supplementary Table S1).

#### 2.1.2 Literature filter based on study type and test organism

In stage 2, we screened the 14297 research articles from stage 1 to select studies based on *in vivo* or *in vitro* experiments in humans or rodents (Figure 1). Here, we have excluded published studies where receptor-based binding assays or *in silico* methods are employed to infer the potential endocrine disruption by a chemical using binding affinity or bioactivity information. Such binding affinity or bioactivity values do not provide sufficient information on whether chemical exposure can actually lead to adverse effects due to endocrine disruption (Baker, 2001). We have also excluded human epidemiological studies due to insufficient mechanistic evidence linking observed adverse effects to potential endocrine disruption upon chemical exposure (Bliatka et al., 2017; Hernandez and Tsatsakis, 2017). The filtration based on study type and test organism led to a subset of 3300 research articles at the end of stage 2 (Supplementary Table S2).

#### 2.1.3 Compilation of tested chemicals from the filtered research articles

In stage 3, we gathered the set of chemicals tested for potential endocrine disruption in any of the 3300 research articles from stage 2. Moreover, we also gathered information on the 2D structure of each tested chemical using PubChem (Kim et al., 2016) and Chemical Abstracts Service (CAS; https://www.cas.org/) databases (Figure 1). Note that we have omitted any tested chemical in the 3300 research articles which could not be mapped to its 2D structure using standard chemical databases. At the end of stage 3, we compiled 1626 chemicals along with their 2D structures that were tested for endocrine disruption in humans or rodents in at least one of the filtered research articles from stage 2 (Supplementary Table S3).

#### 2.1.4 Identification of potential EDCs with supporting evidence for systems-level endocrine-mediated perturbations

In stage 4, we identify potential EDCs among the 1626 chemicals compiled in stage 3 by assessing the significance of observed effects for endocrine disruption upon exposure in published experiments in humans or rodents (Figure 1).

Prior to this assessment of supporting evidence for endocrine disruption upon chemical exposure, we excluded a tested chemical or its published experiment based on the following criteria (Figure 1):

a. Chemical is a natural hormone.
b. Chemical was tested as part of a mixture in the published experiment. This criterion reflects our choice to include chemicals which as single entities can cause endocrine disruption upon exposure.
c. Chemical was tested for therapeutic relevance in the published experiment.

Moreover, we excluded published experiments which contain evidence for endocrine disruption upon chemical exposure in an *in vitro* rodent system. Since the observed effects in an *in vitro* rodent system do not adequately reflect the complexities observed in humans, the last criterion omits such evidence in the published literature (Figure 1). For the next phase of the workflow, we filtered chemicals and their associated literature which pass the above-mentioned criteria.

For each chemical which passed the above-mentioned criteria, we next evaluated the level of supporting evidence for endocrine disruption in humans or rodents upon exposure based on published experiments contained in the filtered research articles. For this evaluation, we manually compiled the *observed effects* upon exposure of each chemical in associated published experiments in humans or rodents. A published experiment in humans or rodents is considered as *strong supporting evidence* for endocrine disruption by a chemical if the chemical upon exposure leads to *observed effects or endpoints* related to endocrine-specific perturbations such as changes in morphology, physiology, growth, reproduction, development and lifespan (WHO/UNEP, 2013). Thereafter, if a chemical has at least one published experiment with strong supporting evidence for endocrine disruption upon exposure, then it is identified as a *potential EDC* in stage 4 of the workflow. At the end of stage 4, we identified 686 potential EDCs with supporting evidence of endocrine-mediated perturbations in published literature spanning 1796 research articles (Supplementary Table S4).

### 2.2 Compilation of endocrine-mediated endpoints and their classification into systems-level perturbations

For the identification of EDCs, we have manually compiled the *observed effects or endpoints* related to endocrine-specific perturbations reported in published experiments on chemical exposure in humans or rodents (Figure 1). This compiled list of observed effects or endpoints was then used to assess the level of supporting evidence for endocrine disruption upon chemical exposure. In order to standardize the reported evidence for an EDC, we undertook an extensive manual effort to unify the biological terms used to describe the observed effects or endpoints related to endocrine-specific perturbations in published experiments upon chemical exposure. This standardization effort led to a comprehensive list of 514 *endocrine-mediated endpoints* which refer to the *adverse effects* such as changes in morphology, physiology, growth, reproduction, development and lifespan that may be observed in experiments after the administration or ingestion of a tested chemical (Supplementary Table S5). For the 686 EDCs, we have also compiled the observed adverse effects in terms of these 514 endocrine-mediated endpoints from published experiments in supporting literature.

EDCs perturb the normal functioning of the human endocrine system which consists of several glands that secrete hormones which in turn regulate diverse biological functions such as development, growth, reproduction, metabolism, immunity and behaviour. Hence, exposure to EDCs can have adverse effects in several biological processes regulated by the human endocrine system (Figure 2). In addition, the endocrine-related processes perturbed by EDCs can also induce cancer in humans (Diamanti-Kandarakis et al., 2009; Gore et al., 2015; WHO/UNEP, 2013). Motivated by the major biological processes controlled by the human endocrine system, we have classified the 514 endocrine-mediated endpoints into 7 systems-level perturbations, namely,

a. Reproductive endocrine-mediated perturbations (RT),
b. Developmental endocrine-mediated perturbations (DT),
c. Metabolic endocrine-mediated perturbations (MT),
d. Immunological endocrine-mediated perturbations (IT),
e. Neurological endocrine-mediated perturbations (NT),
f. Hepatic endocrine-mediated perturbations (HT), and
g. Endocrine-mediated cancer (CT).

In Supplementary Table S5, we list the 514 endocrine-mediated endpoints and their categorization into 7 systems-level endocrine-mediated perturbations.

We highlight that future studies and toxicological databases can leverage our comprehensive list of 514 endocrine-mediated endpoints and their categorization into 7 systems-level perturbations while reporting or documenting the adverse effects related to endocrine disruption from experiments related to chemical exposure. Hence, this work also contributes towards development of a unified biological vocabulary to describe toxicity profiles of chemicals.

### 2.3 Compilation of dosage information for observed endocrine-mediated endpoints

In stage 4 of the workflow, we have also compiled the dosage values for each EDC at which the endocrine-mediated endpoints are observed in the published experiments (Figure 1).

Firstly, we have gathered the test dosage values for each EDC in appropriate units from the published experiments. Secondly, we have identified the effective dosage value among the test dosage values at which a particular endocrine-mediated endpoint is observed upon EDC exposure in the published experiment. Thirdly, the published experiments with supporting evidence for endocrine disruption by EDCs employ different units to report the test and effective dosage values. Thus, we undertook a significant effort to convert and express the test and effective dosage values taken from published experiments on EDCs in a uniform format wherever possible. Based on this effort, we realized that the different units used to report the test and effective dosage values of EDCs in published experiments can be classified into two broad categories:

a. *Dose* which gives the amount of chemical that is administered directly to the test organism in the experiment.
b. *Concentration* which gives the amount of chemical present in another substance such as food, soil or water that is administered to the test organism in the experiment.

Moreover, only a fraction of the published experiments on EDCs report dosage values normalized by the body weight of the individual test organism and duration of exposure. For example, if a published experiment on EDC reports the dosage value in the unit mg/kg/day then this gives the amount of chemical administered per kg of the body weight of the test organism per day. Due to the above-mentioned limitations, we were able to convert the different units used in published experiments to report the dosage values of EDCs into 19 standardized units. Supplementary Table S6 lists these 19 standardized units which were used to compile the dosage values of EDCs specific to endocrine-mediated endpoints from published experiments. For each EDC, we have compiled the test and effective dosage values specific to endocrine-mediated endpoints in standardized units, and this information is readily available via the DEDuCT webserver.

### 2.4 NOAEL and LOAEL information for EDCs

Natural hormones in human body can carry out their physiological functions at very low concentration. EDCs are known to interfere with the endocrine system by mimicking the natural hormones. Thus, it is important for risk assessment of EDCs to understand the adverse effects caused by their low dose exposure (Vandenberg, 2014; Vandenberg et al., 2012; Welshons et al., 2003). In this direction, our compilation of the test and effective dosage values for EDCs from published experiments can be leveraged to elucidate such low dose effects.

Specifically, we have used the test and effective dosage values for EDCs in DEDuCT to determine the following dose-response measures:

a. *No Observed Adverse Effect Level* (*NOAEL*) gives the highest dose of an EDC at which no observed effects or endocrine-mediated endpoints are seen in the published experiments.
b. *Low Observed Adverse Effect Level* (*LOAEL*) gives the lowest dose of an EDC at which any one of the observed effects or endocrine-mediated endpoints are seen in the published experiments.

Note that the supporting evidence for the EDCs in DEDuCT has been compiled from three different types of published experiments, namely, *in vivo* or *in vitro* experiments in humans or *in vivo* experiments in rodents. In cases where the supporting evidence for an EDC comes from more than one type of published experiment, we determine the NOAEL and LOAEL values for the EDC separately for different types of published experiments (Supplementary Table S7). Moreover, the supporting evidence for an EDC in DEDuCT may come from published experiments employing different units to specify test and effective dosage values as discussed in the last section. In such cases, we determine the NOAEL and LOAEL values for the EDC separately for different standardized units across the published experiments (Supplementary Table S7). Note that we did not compile the information on the route and duration of EDC exposure from published experiments in DEDuCT. Supplementary Table S7 lists the NOAEL and LOAEL values for EDCs in DEDuCT.

### 2.5 Classification of EDCs

#### 2.5.1 Based on the type of supporting evidence in published experiments

We have classified the 686 EDCs in DEDuCT into 4 categories based on the type of supporting evidence in published experiments. EDCs in category I have supporting evidence from *in vivo* human experiments, category II from *in vivo* rodent and *in vitro* human experiments but not from *in vivo* human experiments, category III from only *in vivo* rodent experiments, and category IV from only *in vitro* human experiments (Supplementary Table S8). Thus, potential EDCs in category I have the highest level of supporting evidence in published experiments followed by category II, III and IV, respectively.

#### 2.5.2 Based on the environmental source

Based on the environmental source of EDCs, we have classified the 686 EDCs into 7 broad categories, namely, Agricultural and farming, Consumer products, Industry, Intermediates, Medicine and health care, Natural sources, and Pollutant (Figure 3). Furthermore, the 7 broad categories of EDCs were further classified into 48 sub-categories (Figure 3). Note that this environmental source-based classification of EDCs is overlapping, that is, a given EDC may belong to multiple broad or sub-categories.

#### 2.5.3 Based on chemical structure

We have employed the web-based application ClassyFire (Djoumbou Feunang et al., 2016) (http://classyfire.wishartlab.com/) to obtain a chemical classification of the 686 EDCs. Note that ClassyFire gives a non-overlapping hierarchical chemical classification based on the structure and composition of the molecule. Using ClassyFire, the 686 EDCs were classified into two chemical kingdoms, namely, organic and inorganic compounds, respectively (Figure 4). Moreover, the EDCs in the organic kingdom can be further classified into 19 super-classes while those in the inorganic kingdom fall into 3 super-classes (Figure 4).

### 2.6 Physicochemical properties and molecular descriptors

For the 686 EDCs, we obtained the 2D chemical structure from Pubchem and CAS databases. Thereafter, Balloon (Vainio and Johnson, 2007) (http://users.abo.fi/mivainio/balloon/) and Open Babel (O’Boyle et al., 2011) (http://openbabel.org/) with Merck Molecular Force Field (MMFF94) were used to generate the lowest energy 3D structure of the EDCs. RDKit (https://www.rdkit.org/) was used to compute the basic physicochemical properties of the EDCs. In addition, we have also computed the one-dimensional (1D), 2D and 3D molecular descriptors using PaDEL (Yap, 2011) (http://www.yapcwsoft.com/dd/padeldescriptor/), RDKit and Pybel (O’Boyle et al., 2008). For each EDC, PaDEL, RDKit and Pybel gave 1875, 213 and 14 descriptors, respectively. For each EDC, we have made its 2D and 3D chemical structure, physicochemical properties and molecular descriptors readily available via the DEDuCT webserver, and this information can aid future efforts to develop computational toxicity models based on structure-activity relationships.

### 2.7 Chemical similarity network

We constructed the chemical similarity network (CSN) of the 686 EDCs as follows. In the CSN, nodes are EDCs and the edge weights reflect the extent of chemical similarity between pairs of EDCs. To assign the edge weight or to quantify the chemical similarity between two EDCs, we computed the Tanimoto coefficient (Bajusz et al., 2015; Tanimoto, 1957) between two EDCs using the extended connectivity fingerprints (ECFP4) (Rogers and Hahn, 2010) by applying Morgan algorithm (Morgan, 1965) as implemented in RDKit. Since Tanimoto coefficient for any pair of chemicals is in the range 0 to 1, the edge weights in the CSN are in the same range.

To visualize the high similarity backbone of the CSN, we decided to omit edges with weights below a chosen threshold value signifying poor chemical similarity. Rather than choosing an arbitrary threshold value to construct this high CSN, we have investigated the size of the largest connected component of the CSN as a function of the increasing threshold value for omitting edges (Supplementary Figure S1A). Note that the size of the largest connected component reflects the overall connectivity of the network. Based on this investigation, we find that there is a sharp decrease in the size of the largest connected component of the CSN obtained after omitting edges below the threshold value of 0.450 (Supplementary Figure S1A). Subsequently, we used this threshold value of 0.450 to construct the high CSN of the 686 EDCs (Figure 5A; Supplementary Table S9).

### 2.8 Target genes of EDCs based on ToxCast assays

Information on the target genes of EDCs can elucidate molecular initiating events leading to adverse effects upon chemical exposure. ToxCast (Dix et al., 2006) uses high-throughput assays designed to screen toxic chemicals based on perturbation of biological activities upon exposure. To date, ToxCast has screened more than 9000 chemicals using more than 900 high-throughput assays. We used the ToxCast invitroDB3 dataset released in October 2018 (Toxicology, 2018) to obtain the list of perturbed genes upon EDC exposure.

The assay summary information file (Assay_Summary_180918.csv) contains the detailed annotation of the ToxCast assays including assay type, assay component, assay component endpoint, assay target information, cell lines used for the assay, and assay citation. Using the assay component endpoint of a ToxCast assay, one can obtain the observed biological effect such as changes in gene expression upon chemical exposure. In practice, the assay component endpoint of a ToxCast assay may correspond to one or more target genes. The assay activity information file (hitc_Matrix_180918.csv) provides a list of *active* or *inactive* chemicals based on the potency of the chemical to produce a significant biological effect captured via 1504 assay component endpoints of different ToxCast assays. In this work, we restrict to ToxCast assays and their corresponding assay component endpoints that are specific to human. If a tested chemical is *active* for a particular assay component endpoint of a ToxCast assay, then the corresponding *gene* is assigned to be the *target* of the chemical.

Of the 686 potential EDCs in DEDuCT, we found target genes for 383 EDCs based on 1228 ToxCast assay component endpoints specific to human. Supplementary Table S10 gives the target genes of these 383 EDCs based on ToxCast assay component endpoints specific to human. We remark that it is possible to expand this information on target genes of EDCs using toxicological databases such as CTD (Mattingly et al., 2003), however, CTD compiles target information from both experiments and computational predictions.

### 2.9 Target similarity network

For the 383 EDCs with information on target genes from ToxCast assays, we have constructed a target similarity network (TSN) based on shared target genes between pairs of EDCs. In the TSN, nodes are EDCs and edge weights signify the target similarity between pairs of EDCs. To quantify the similarity between two sets of target genes corresponding to a pair of EDCs, we use the standard measure, Jaccard index (Jaccard, 1912), given by the ratio of the number of elements in the intersection over the number of elements in the union of the two sets of target genes. By construction, Jaccard index is in the range 0 to 1. Jaccard index between two EDCs is 0 if they have no target genes in common, and it is 1 if they have all target genes in common.

To visualize the high similarity backbone of the TSN, we decided to omit edges with weights below a chosen Jaccard index value signifying poor target similarity between pairs of EDCs. Rather than choosing an arbitrary Jaccard index value to construct this high TSN, we have investigated the size of the largest connected component of the TSN as a function of the increasing Jaccard index value for omitting edges (Supplementary Figure S1B). Based on this investigation, we find that there is a sharp decrease in the size of the largest connected component of the TSN obtained after omitting edges below the Jaccard index of 0.517 (Supplementary Figure S1B). Subsequently, we used this threshold Jaccard index of 0.517 to construct the high TSN of the 383 EDCs (Figure 5B; Supplementary Table S11).

### 2.10 Predicted ADMET properties

Absorption, Distribution, Metabolism, Excretion and Toxicity (ADMET) properties can be utilized for the toxicity assessment of chemicals. Thus, several computational tools have been developed to predict the ADMET properties of chemicals such as admetSAR 2.0 (Yang et al., 2019), pkCSM (Pires et al., 2015), ProTox (Banerjee et al., 2018), SwissADME (Daina et al., 2017), Toxtree 2.6.1 (Patlewicz et al., 2008) and vNN server (Schyman et al., 2017). We have employed these tools to predict the ADMET properties of the 686 potential EDCs in DEDuCT.

Absorption properties of a chemical reflect its ability to be absorbed from intestine to bloodstream. The predicted absorption properties for EDCs include Caco-2 permeability, human intestinal absorption (HIA), human oral bioavailability and skin permeability (log Kp). Distribution properties of a chemical shed light on its availability in other parts of the body after being absorbed into bloodstream. The predicted distribution properties for EDCs include blood-brain barrier (BBB), CNS permeability, fraction unbound in human, P-glycoprotein inhibitor, P-glycoprotein substrate, plasma protein binding, steady state volume of distribution (VDss) and subcellular localization. Metabolism properties of a chemical describe its conversion into metabolites through enzymatic breakdown prior to elimination from the human body. The predicted metabolism properties for EDCs include assessment to act as a substrate or inhibitor of CYP450 enzymes, human bile salt export pump (BSEP), human liver microsomal (HLM) stability assay, human multidrug and toxin extrusion (MATE) transporter, organic anion-transporting polypeptides (OATP) and UDP-glucuronosyltransferases (UGT) catalysis. The predicted excretion properties for EDCs include total clearance rate and the ability to inhibit or act as a substrate for renal organic cation transporter 2 (OCT2). The predicted toxicological properties for EDCs include biodegradation capacity, carcinogenicity, Cramer’s rule, cytotoxicity, hepatotoxicity, hERG inhibitors, maximum recommended tolerated dose (MRTD), mitochondrial membrane potential (MMP), rat oral toxicity and skin sensitization. Supplementary Table S12 lists the predicted ADMET properties by different tools used here. For each EDC, we have made the predicted ADMET properties readily available via the DEDuCT webserver.

### 2.11 Web interface and database management system

We have created an online resource, **D**atabase of **E**ndocrine **D**isr**u**pting **C**hemicals and their **T**oxicity profiles (DEDuCT), which contains detailed information on the 686 potential EDCs with supporting evidence compiled from 1796 published research articles. Importantly, DEDuCT compiles the above-mentioned information on the 686 EDCs such as the endocrine-mediated endpoints, systems-level endocrine-mediated perturbations, dosage value specific to endpoints, type of supporting evidence based classification, environmental source-based classification, 2D and 3D chemical structures, chemical classification, physicochemical properties, molecular descriptors, predicted ADMET properties and target genes. DEDuCT is accessible at: https://cb.imsc.res.in/deduct/.

The web interface of DEDuCT was created using PHP (http://php.net/), HTML, CSS, Bootstrap 4, and jQuery (https://jquery.com/). To facilitate interactive visualization, we have used Google Charts (https://developers.google.com/chart/), D3.js (https://d3js.org/), Cytoscape.js (http://js.cytoscape.org/) and JSmol (http://jmol.sourceforge.net/) in the web interface. The compiled database on EDCs is stored using MariaDB (https://mariadb.org/), and the information from the database is retrieved using Structured Query Language (SQL). DEDuCT website is hosted on Apache (https://httpd.apache.org/) webserver running on Debian 9.4 Linux Operating System.

## 3. Results and Discussion

### 3.1 DEDuCT – a curated knowledgebase on EDCs with supporting evidence from published experiments in humans or rodents

Using the detailed workflow shown in Figure 1, we have compiled 686 potential EDCs with supporting evidence of endocrine disruption upon exposure in experiments on humans or rodents from 1796 published research articles (Methods). Our webserver, **D**atabase of **E**ndocrine **D**isr**u**pting **C**hemicals and their **T**oxicity profiles (DEDuCT), contains the compiled information on these 686 potential EDCs (Methods).

We have classified the 686 EDCs into 4 categories (I-IV) based on the type of supporting evidence in published experiments for endocrine disruption (Methods; Supplementary Table S8). Of the 686 EDCs, 7, 142, 367 and 170 are in category I, II, III and IV, respectively (Supplementary Table S8). These 142, 367 and 170 potential EDCs in categories II, III and IV, respectively, require additional experimentation and further risk assessment for their potential risk to humankind.

We have also classified the 686 EDCs into 7 broad categories and 48 sub-categories based on their environmental source (Methods; Figure 3). Majority of EDCs in DEDuCT are used in consumer products (Figure 3). Moreover, we also provide a hierarchical classification of the 686 EDCs into chemical kingdoms and chemical super-classes based on their chemical structure (Methods; Figure 4). Of the 686 EDCs, 644 are organic and 42 are inorganic (Figure 4). Among the 644 organic EDCs, the largest fraction belongs to the chemical super-class Benzenoids (Figure 4).

As supporting evidence for the identification of EDCs, we have compiled the observed endocrine-specific effects upon chemical exposure from published experiments in the form of 514 endocrine-mediated endpoints (Methods; Figure 2). Moreover, the 514 endocrine-mediated endpoints were further classified into 7 systems-level perturbations based on the affected biological processes controlled by the endocrine system (Methods; Figure 2). In Figure 6A, we show the occurrence of these 7 systems-level perturbations in the supporting published experiments for the 686 EDCs. Among the 686 EDCs, it is seen that 535 have supporting evidence for reproductive perturbations and 315 for metabolic perturbations (Figure 6A). Thus, majority of EDCs in DEDuCT have supporting evidence for adverse effects on the reproductive system followed by metabolism.

**Figure 6:**
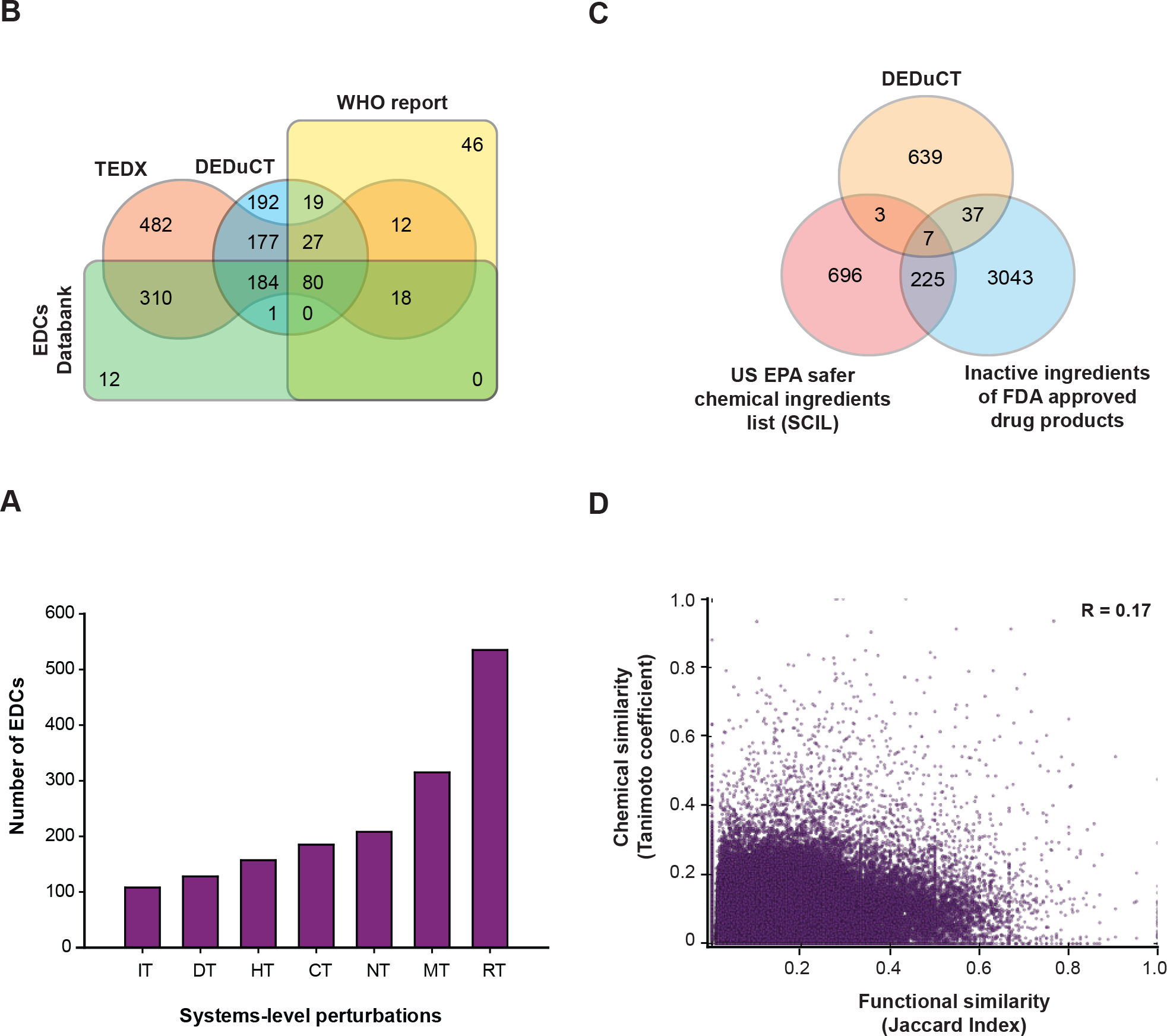
(**A)** Histogram shows the occurrence of 7 systems-level perturbations in the supporting evidence compiled from published experiments for the 686 EDCs. Majority of EDCs in DEDuCT have adverse effects on the reproductive system followed by metabolism. **(B)** Comparison of the 686 EDCs in DEDuCT with those in the WHO report, TEDX and EDCs Databank. From the Venn diagram, it is seen that 198 EDCs in DEDuCT are not captured in the three other existing resources. **(C)** Comparison of the 686 EDCs in DEDuCT with the US EPA safer chemical ingredient list (SCIL) and the list of inactive ingredients of FDA approved drug products. 10 EDCs are present in the SCIL while 44 EDCs are present in FDA inactive ingredients list. **(D)** Scatter plot of functional similarity between sets of target genes for pairs of EDCs as a function of chemical structure similarity between pairs of EDCs. We find no significant correlation (Pearson correlation coefficient R=0.17) between the structural and functional similarity of EDCs.

Using the *Browse* section in the web interface of DEDuCT, users can view the EDCs based on their type of supporting evidence or environmental source or chemical classification or systems-level perturbations (Supplementary Figure S2). Using the *Simple search* option in DEDuCT, users can search for individual EDCs using chemical name or standard identifier (Supplementary Figure S2). Using the *Physicochemical filter* option in DEDuCT, users can also filter EDCs based on their physicochemical properties such as molecular weight, number of hydrogen bond donors or acceptors, number of rotatable bonds (Supplementary Figure S2). By clicking the chemical name of any EDC in DEDuCT, users can view the entire compiled information including supporting evidence and dosage information (Methods).

### 3.2 Comparison of DEDuCT with existing resources on EDCs

Besides DEDuCT, there are at least five existing resources on EDCs including the WHO report (WHO/UNEP, 2013), TEDX, EDCs Databank (Montes-Grajales and Olivero-Verbel, 2015), EDKB (Ding et al., 2010) and Endocrine Disruptor Screening Program (EDSP; https://www.epa.gov/endocrine-disruption) of United States Environmental Protection Agency (US EPA).

Both the WHO report and TEDX contain manually curated information on EDCs based on published literature evidence including *in vivo*, *in vitro*, environmental monitoring and epidemiological studies. EDCs Databank compiles EDCs from the TEDX and the EU list of potential endocrine disruptors followed by PubMed search to associate literature evidence with EDCs. Also, EDCs Databank provides detailed information on compiled EDCs including their 2D and 3D structure, physicochemical properties, categorization based on exposure source and external links to other toxicological databases.

In addition to extensive PubMed mining to identify published experiments on EDCs, DEDuCT integrates information from the WHO report, TEDX and EDCs Databank. Since the WHO report, TEDX and EDCs Databank are not limited to *in vivo* or *in vitro* studies in humans and *in vivo* studies in rodents, we have filtered the compiled experimental evidence in the three resources using our workflow to identify potential EDCs with experimental evidence for endocrine disruption in humans or rodents (Methods; Figure 1). Importantly, unlike DEDuCT, the WHO report, TEDX and EDCs Databank do not compile the observed endocrine-mediated endpoints and systems-level perturbations upon EDC exposure (Table 1). Table 1 presents a detailed comparison of DEDuCT with the WHO report, TEDX and EDCs Databank. We find that 198 out of the 686 potential EDCs (28.9%) in DEDuCT are not captured in any of the three existing resources (Figure 6B). Note that we were unable to find supporting evidence for endocrine disruption upon exposure in published experiments on humans or rodents for several chemicals in the WHO report or TEDX or EDCs Databank, and thus, such chemicals are not contained in DEDuCT (Figure 6B; Methods).

**Table 1:**
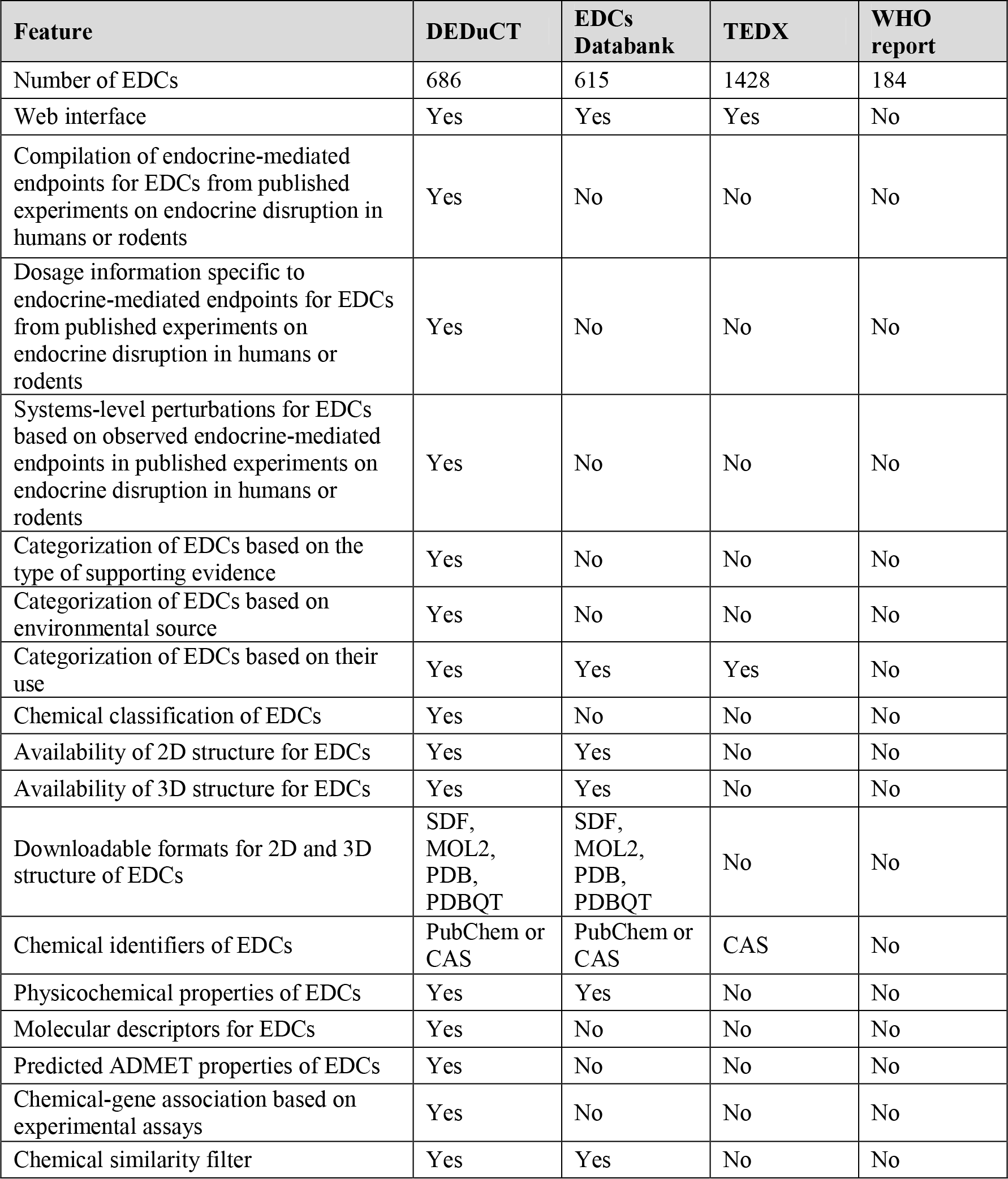
Comparison of the information on EDCs in DEDuCT with three existing resources, namely, EDCs Databank, TEDX and WHO report.

In this work, we decided to not include information contained in two existing resources, EDKB (Ding et al., 2010) and EDSP due to the following reasons. EDKB compiles EDCs based on multiple receptor binding assays and *in silico* QSAR studies, and such evidence is ignored in our workflow to identify EDCs (Methods; Figure 1). Thus, we do not include information in EDKB as it does not contain supporting evidence for EDCs based on observed adverse effects in *in vivo* or *in vitro* experiments in humans or rodents. EDSP of US EPA is a regulatory framework to screen and prioritize chemicals with potential to interact with the endocrine system. EDSP has carried out several hormonal assays in test organisms such as human, rat, fish and amphibians to determine the potency of a chemical to interact with the endocrine system. EDSP identifies a chemical to be an EDC if the chemical displays consistent evidence of endocrine disruption across all hormonal assays carried out by them. As highlighted by Zoeller et al. (Zoeller et al., 2012), the weight of evidence used by EDSP to identify EDCs is too stringent which leads to omission of several chemicals with significant endocrine-specific effects. Specifically, in the EDSP Tier 1 screening of 52 chemicals, 18 were determined to have conclusive evidence for endocrine disruption while 34 have inconclusive evidence according to EDSP. However, a closer inspection of the 34 chemicals determined by EDSP to have inconclusive evidence finds well known EDCs such as Chlorpyrifos and 2,4-Dichlorophenoxyacetic acid highlighted by the WHO report and the Endocrine society (https://www.endocrine.org/topics/edc). Thus, we decided not to include information from EDSP in DEDuCT.

We remark that the primary goal of DEDuCT is to create a comprehensive curated compilation of published experiments on endocrine disruption in humans or rodents upon chemical exposure. DEDuCT was built with the sole intention to enable basic research towards understanding of systems-level perturbations upon EDC exposure.

### 3.3 Comparison with US EPA safer chemical ingredients list

US EPA has evaluated and released a safer chemical ingredients list (SCIL) of 931 chemicals based on their functional use categories as part of its safer choice program (https://www.epa.gov/saferchoice/safer-ingredients). In SCIL, US EPA has labelled chemicals of low concern by *green circle*, chemicals of low concern for which additional data is required by *green half-circle*, chemicals satisfying safer choice criteria only for a particular functional use while possibly displaying hazardous profile in other uses by *yellow triangle*, and chemicals unsuitable for use in consumer products by *grey square*. We decided to compare the subset of 930 SCIL chemicals labelled by green circle or green half-circle or yellow triangle with the 686 potential EDCs in DEDuCT. 10 out of the 686 potential EDCs were found in the SCIL (Figure 6C).

None of these 10 potential EDCs in SCIL are listed under category I EDCs in DEDuCT with supporting evidence for endocrine disruption from *in vivo* human experiments. Of these 10 potential EDCs, 1, 7 and 2 are in category II, III and IV, respectively. Benzyl salicylate is the only chemical in SCIL that is listed as category II EDC in DEDuCT with supporting evidence for endocrine disruption from *in vivo* rodent and *in vitro* human experiments while lacking evidence from *in vivo* human experiments. As Benzyl salicylate is labelled by yellow triangle in SCIL based on the functional use category of fragrances, this suggests that this chemical may have potential to display hazardous profile in other use categories. For improved risk assessment, there is need to further evaluate and gather additional evidence for potential EDCs listed in the categories II, III and IV of DEDuCT.

### 3.4 Comparison with inactive ingredients of FDA approved drug products

We have also compared the list of 3312 inactive ingredients used in US Food and Drug Administration (FDA) approved drug products from inactive ingredient database (https://www.accessdata.fda.gov) with 686 potential EDCs in DEDuCT. Inactive ingredients in a drug are the chemicals that do not have any pharmacological effect and these include colorants, drug preservatives and flavouring agents. We find that 44 of the 686 potential EDCs are used as inactive ingredients in FDA approved drugs (Figure 6C). None of these 44 potential EDCs are listed under category I EDCs in DEDuCT. Of 44 potential EDCs in FDA inactive ingredients list, 7 chemicals (Caffeine, Trichloroethylene, Diethyl phthalate, Butyl p-hydroxybenzoate, Methyl p-hydroxybenzoate, Ethyl p-hydroxybenzoate, Butylated hydroxyanisole) are in category II, 30 in category III, and 7 in category IV of DEDuCT. For better risk assessment, these 44 potential EDCs in FDA inactive ingredients list require additional evidence from *in vivo* human experiments considering the effective dosage, route of exposure, and duration of exposure.

### 3.5 EDCs differ both in their chemical structure and set of target genes

We investigated the extent of chemical similarity between the 686 EDCs in DEDuCT. For this, we constructed the high chemical similarity network (CSN) for the 686 EDCs shown in Figure 5A (Methods; Supplementary Table S9). In Figure 5A, it is seen that the high CSN has a large connected component of 255 EDCs, many small components of 2 or more EDCs and many isolated EDCs. From this analysis, we concluded that EDCs can be dissimilar in their chemical structure.

We next investigated the similarity between the target genes of EDCs in our resource. Based on ToxCast assays, we were able to obtain the set of target genes for 383 out of 686 EDCs (Methods; Supplementary Table S10). Thereafter, we constructed the high target similarity network (TSN) for the 383 EDCs shown in Figure 5B (Methods; Supplementary Table S11). In Figure 5B, it is seen that the high TSN has a large connected component of 199 EDCs, many small components of 2 or more EDCs and many isolated EDCs. Based on the limited information on target genes from ToxCast assays, we concluded that EDCs can have very different set of target genes.

Lastly, we investigated whether there is any relationship between structural similarity and functional similarity of EDCs. Recall that we have quantified the structural similarity between two EDCs using the Tanimoto coefficient and the functional similarity or commonality between the sets of target genes for two EDCs using the Jaccard index (Methods). In Figure 6D, we plot this structural similarity between pairs of EDCs as a function of the functional similarity between sets of target genes for pairs of EDCs. Based on the limited information on target genes for 383 EDCs from ToxCast assays, we find no significant correlation between structural similarity and functional similarity of EDCs (Figure 6D).

These observations underscore the challenge in developing computational models to predict EDCs. Since traditional computational toxicity models based on quantitative structure activity relationship (QSAR) use chemical similarity and bioactivity information for their predictions, our results based on high CSN and high TSN suggest that such models for EDCs are unlikely to have high predictive power. Alternatively, computational systems toxicity models leveraging information in DEDuCT on chemical structure, dosage information, set of target genes and systems-level perturbations of EDCs may have better predictive power.

### 3.6 Evaluation of the sensitivity of toxicity predictors using compiled experimental evidence in DEDuCT

Several computational toxicity predictors such as admetSAR 2.0 (Yang et al., 2019), pkCSM (Pires et al., 2015), ProTox (Banerjee et al., 2018), SwissADME (Daina et al., 2017), Toxtree 2.6.1 (Patlewicz et al., 2008) and vNN server (Schyman et al., 2017) have been developed for risk assessment of chemicals. We have used these tools to predict the ADMET properties of the 686 EDCs, and this information is readily available from DEDuCT webserver (Methods; Supplementary Table S12). Since DEDuCT compiles experimentally observed toxicity profiles or endocrine-mediated endpoints for the 686 EDCs from supporting literature, we decided to utilise this compiled experimental evidence as a positive dataset to evaluate the *sensitivity* of computational toxicity prediction tools.

In DEDuCT, 157 EDCs have experimental evidence to cause hepatic endocrine-mediated perturbations. Among the toxicity predictors, admetSAR 2.0, pkCSM and vNN server can predict the hepatotoxicity of chemicals. Of these 157 EDCs, admetSAR 2.0, pkCSM and vNN server gave correct prediction for 60, 22 and 41 EDCs, respectively. Thus, the sensitivity for predicting hepatotoxicity of EDCs by admetSAR 2.0, pkCSM and vNN server are 0.39, 0.14 and 0.26, respectively, based on our dataset.

In DEDuCT, 185 EDCs have experimental evidence to cause endocrine-mediated cancer. Among the toxicity predictors, admetSAR 2.0 and Toxtree 2.6.1 can predict the carcinogenicity of chemicals. Of these 185 EDCs, admetSAR 2.0 predicted 56 while Toxtree 2.6.1 predicted none to be carcinogens. Thus, the sensitivity for predicting carcinogenicity of EDCs by admetSAR 2.0 and Toxtree 2.6.1 is 0.30 and 0.0, respectively, based on our dataset.

admetSAR 2.0 predicted 127 out of the 185 EDCs with experimental evidence to cause cancer in DEDuCT to be non-carcinogens, and we have compared these 127 EDCs with the potential carcinogens released by the International Agency for Research on Cancer (IARC) Monographs (Loomis et al., 2018) (https://monographs.iarc.fr/) and the Report on Carcinogens (RoC) by the National Toxicology Program (https://ntp.niehs.nih.gov/pubhealth/roc/). Based on this comparison, we found 9 of the 127 EDCs predicted as non-carcinogens by admetSAR 2.0 were listed as potential carcinogens in IARC Monographs and RoC. Notably, 3 of the 127 EDCs, namely, benzo[a]pyrene, diethylstilbesterol and pentachlorophenol are categorized as group 1 potential carcinogens for human by IARC Monographs.

In summary, this evaluation of the computational toxicity tools for prediction of hepatotoxicity and carcinogenicity of EDCs based on the compiled experimental evidence in DEDuCT suggests lack of significant predictive power. A possible interim solution towards increasing the predictive power of the existing tools will be to update their positive training dataset with experimental information on EDCs from DEDuCT.

## 4. Conclusions

EDCs are a group of chemicals of emerging concern which are omnipresent in our environment. Since endocrine disruption mechanism is a special form of toxicity, the risk assessment and identification of EDCs remains challenging (WHO/UNEP, 2013). In this work, we have developed a detailed workflow which was employed to identify 686 potential EDCs with supporting evidence for endocrine disruption from published experiments in humans or rodents. Our workflow for identification of EDCs consists of four stages (Figure 1). Firstly, we used text mining and existing resources to compile more than 16000 published research articles with potential information on EDCs. Secondly, we manually inspected these published research articles to filter a subset of 3300 articles containing published experiments for endocrine disruption in humans or rodents upon chemical exposure. Thirdly, we compiled the list of 1626 chemicals tested for endocrine disruption in at least one of these 3300 research articles. Fourthly, we evaluated the significance of the observed effects in the published experiments for the 1626 tested chemicals for endocrine-specific perturbations, and this led to the identification of 686 potential EDCs with supporting evidence from 1796 published research articles. Importantly, we have compiled, unified and standardized the observed adverse effects upon EDC exposure in published experiments into 514 unique endocrine-mediated endpoints which were further classified into 7 systems-level perturbations (Figure 2). Vitally, we have also compiled the dosage information at which endocrine-mediated endpoints were observed upon EDC exposure in published experiments. Thereafter, we have classified the potential EDCs based on the type of supporting evidence, their environmental source (Figure 3) and their chemical classification (Figure 4). Lastly, we have obtained additional information for the 686 potential EDCs including 2D and 3D chemical structures, physicochemical properties, molecular descriptors, predicted ADMET properties and target genes. DEDuCT webserver contains the entire compiled information on the 686 potential EDCs and is accessible at: https://cb.imsc.res.in/deduct/.

After compiling the 686 potential EDCs, we have compared the US EPA safer chemical ingredients list (SCIL) and FDA inactive ingredients list with DEDuCT. Of the 686 potential EDCs, 10 and 44 are present in SCIL and FDA inactive ingredients list, respectively. However, none of these potential EDCs in SCIL or FDA inactive ingredients list are among category I EDCs in DEDuCT with supporting evidence for endocrine disruption from *in vivo* human experiments. The potential EDCs in SCIL or FDA inactive ingredients list which are in category II, III or IV in DEDuCT require further evaluation of their potential risk based on additional evidence from *in vivo* human experiments.

Furthermore, we have employed a network-centric approach to understand the link between the chemical space of EDCs and their biological target space. Here, we have constructed and analysed two different networks of EDCs, namely, the chemical similarity network (CSN) and the target similarity network (TSN). Based on CSN, we infer that EDCs can be dissimilar in their chemical structure. Based on TSN, we infer that EDCs can have very different set of target genes. Subsequent investigation of the relationship between the chemical structure and biological (gene) targets of EDCs found no correlation. These results in addition to a previous report (Diamanti-Kandarakis et al., 2009) underscore that EDCs can be structurally dissimilar, and this raises potential challenges in developing structure-based predictive computational models for EDCs. Lastly, the compiled experimental evidence for EDCs in DEDuCT was used to evaluate the predictive power of existing computational toxicity tools. Such an evaluation using our compiled dataset suggests that the existing tools for predicting hepatotoxicity and carcinogenicity of chemicals lack significant predictive power. In near future, toxicity predictors can integrate experimental evidence from DEDuCT to improve their predictive power.

An important aspect of EDCs is their ability to exert adverse effects even at low dosage values (Vandenberg, 2014; Vandenberg et al., 2012; Welshons et al., 2003). Our compilation of dosage information at which endocrine-mediated endpoints were observed in published experiments upon individual EDC exposure will further help researchers to understand the low dose exposure effects of EDCs. Also, our large-scale compilation of the observed effects or endpoints along with the systems-level perturbations upon EDC exposure can be visualized as a tripartite network with nodes as EDCs, endocrine-mediated endpoints and systems-level perturbations. Future exploration of this tripartite network will enhance systems-level understanding of perturbed biological pathways upon EDC exposure.

## Acknowledgements

We thank Sanjay Jain for discussions and P. Mangalapandi for technical help in setting up the webserver. AS is supported by intramural funds from the Department of Atomic Energy (DAE) India and a Ramanujan fellowship (SB/S2/RJN-006/2014) from the Science and Engineering Research Board (SERB), Department of Science and Technology (DST) India. The funders have no role in study design, data collection, data analysis, manuscript preparation or decision to publish.

## Author contributions

B.S.K. and J.R. contributed equally to this work and should be considered as Joint-First authors. K.M. and R.P.V. also contributed equally to this work and should be considered as Joint-Second authors. A.S., B.S.K. and J.R. designed research. J.R. and B.S.K. compiled and curated the data from various sources. K.M. designed the webserver. R.P.V. performed the cheminformatic analysis. J.R., K.M. and A.S. prepared the figures and tables. A.S., J.R. and B.S.K. wrote the manuscript. A.S. planned and supervised the project. All authors have read and approved the manuscript.

## Declaration of Interests

Some of the compiled data in this work was submitted for protection to the copyright office, Government of India. Based on this copyright application, the authors were granted a literary copyright (L-79979/2019) by the Government of India and the copyright owner is the authors’ institution, The Institute of Mathematical Sciences, Chennai, India.

## Supplementary Tables

**Supplementary Table S1:** Filtered list of research articles compiled from PubMed query, the WHO report, TEDX and EDCs Databank that contain keywords related to EDCs. This list of 14297 research articles was obtained at the end of stage 1 of the workflow shown in Figure 1.

**Supplementary Table S2:** Filtered list of research articles on EDCs based on study type and test organism. This list of 3300 research articles was obtained at the end of stage 2 of the workflow shown in Figure 1.

**Supplementary Table S3:** List of 1626 chemicals along with their Pubchem and CAS identifiers that were tested for endocrine disruption in humans or rodents in at least one of the filtered research articles from stage 2 of the workflow shown in Figure 1.

**Supplementary Table S4:** Final list of 686 EDCs with supporting evidence of systems-level endocrine-mediated perturbations in published experiments from 1796 research articles.

**Supplementary Table S5:** List of 514 endocrine-mediated endpoints and their categorization into 7 systems-level endocrine-mediated perturbations. List of abbreviations: RT - Reproductive endocrine-mediated perturbations; DT - Developmental endocrine-mediated perturbations; MT - Metabolic endocrine-mediated perturbations; IT - Immunological endocrine-mediated perturbations; NT - Neurological endocrine-mediated perturbations; HT - Hepatic endocrine-mediated perturbations; CT - Endocrine-mediated cancer.

**Supplementary Table S6:** List of standardized units along with their description that were used to compile the dosage values of EDCs specific to endocrine-mediated endpoints from published experiments.

**Supplementary Table S7:** Final list of No observed adverse effect level (NOAEL) and Low observed adverse effect level (LOAEL) values for EDCs based on test and effective dosage values reported in published experiments in supporting literature. Note that the supporting evidence for the EDCs in our resource has been compiled from three different types of published experiments or study types, namely, *in vivo* (IVH) or *in vitro* (IVTH) experiments in humans or *in vivo* (IVR) experiments in rodents. Importantly, NOAEL value for an EDC could not be determined from a published experiment if an endocrine-mediated endpoint or adverse effect is observed at every sampled test dosage, and in such a case, it is only possible to determine the LOAEL value for the EDC.

**Supplementary Table S8:** Classification of the 686 EDCs into 4 categories (I-IV) based on the type of supporting evidence for endocrine disruption in published experiments.

**Supplementary Table S9:** High chemical similarity network (CSN) of the 686 EDCs obtained using the threshold value of 0.450 for the Tanimoto coefficient. Note that the Tanimoto coefficient is a measure used to quantify the chemical similarity between any pair of chemicals. The table lists the edges between pairs of EDCs in the high CSN along with the edge weights given by the Tanimoto coefficient between pairs of EDCs.

**Supplementary Table S10:** List of target genes for 383 EDCs based on 1228 ToxCast assay component endpoints specific to human.

**Supplementary Table S11:** High target similarity network (TSN) of the 383 EDCs obtained using the threshold value of 0.517 for the Jaccard index. Note that the Jaccard index is a measure used to quantify the similarity between the sets of target genes for any pair of EDCs. The table lists the edges between pairs of EDCs in the high TSN along with the edge weights given by the Jaccard index between pairs of EDCs.

**Supplementary Table S12:** The list of ADMET properties predicted by the considered software tools, namely, admetSAR 2.0, pkCSM, ProTox, SwissADME, Toxtree 2.6.1 and vNN server.

## Supplementary Figures

**Supplementary Figure S1:**
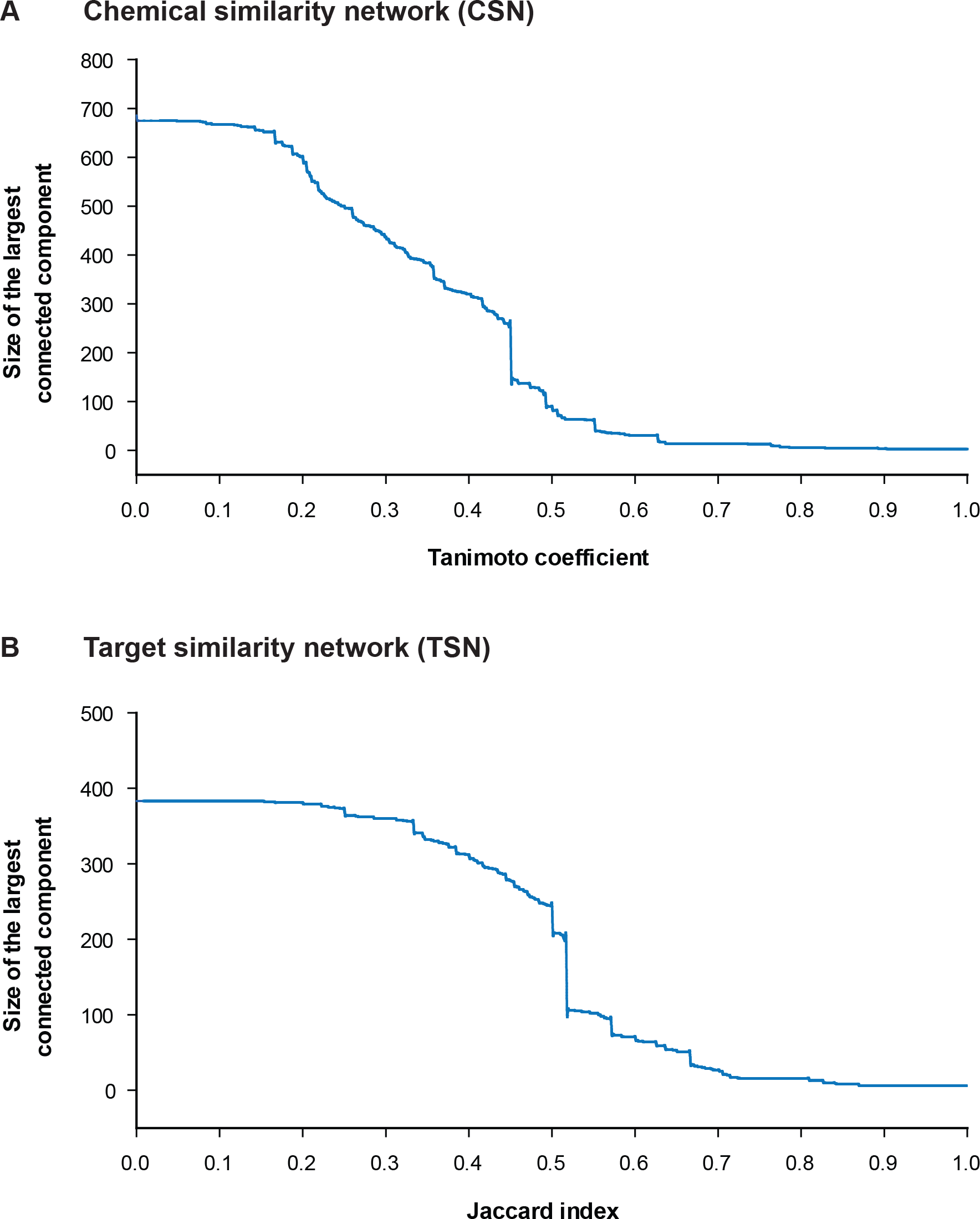
**(A)** The size of the largest connected component of the chemical similarity network (CSN) of EDCs as a function of the increasing Tanimoto coefficient for omitting edges. **(B)** The size of the largest connected component of the target similarity network (TSN) of EDCs as a function of the increasing Jaccard index for omitting edges.

**Supplementary Figure S2:**
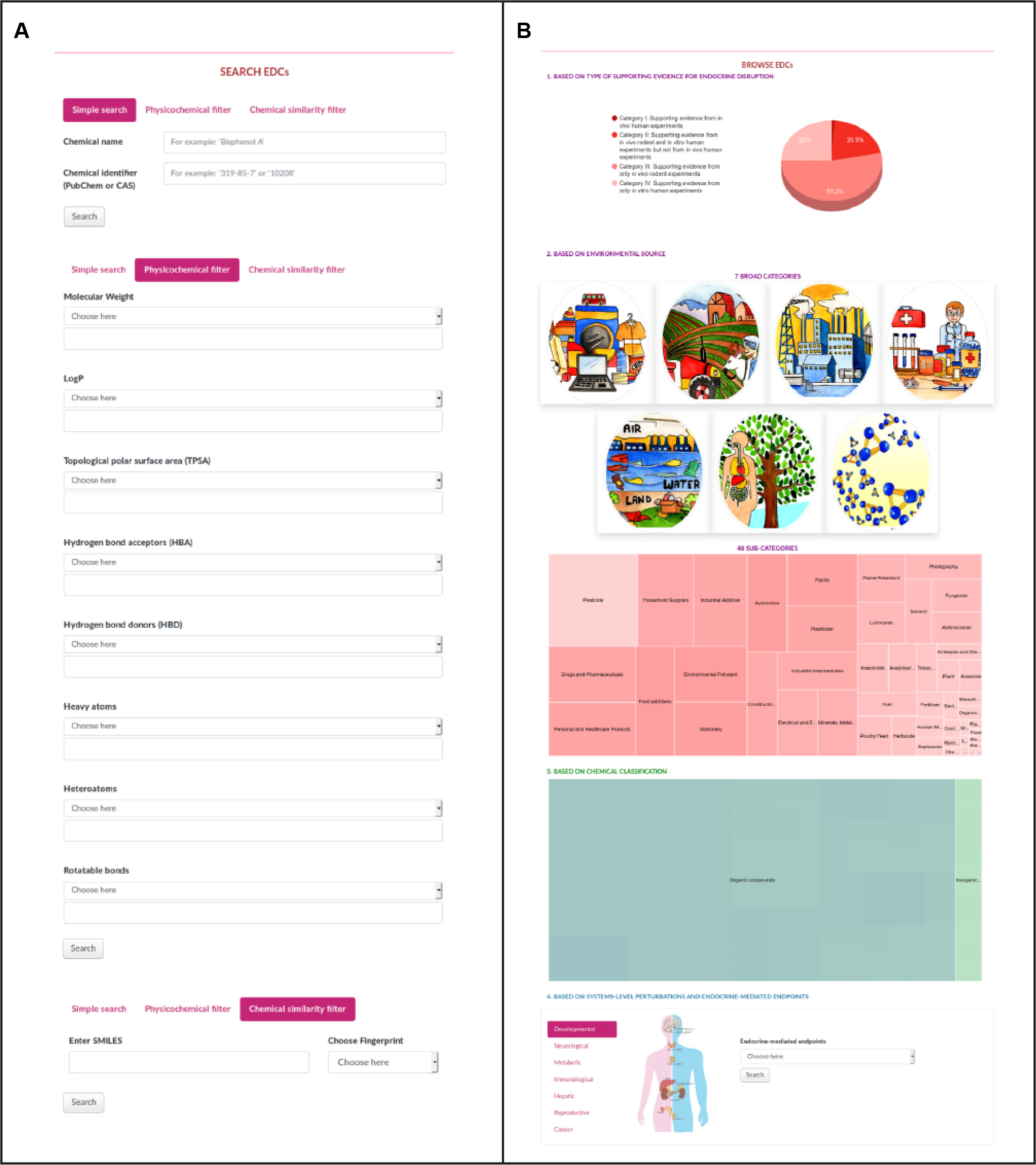
The web interface of DEDuCT. **(A)** The screenshot shows the different search options in our resource to obtain information on EDCs. Simple search option in DEDuCT can be used to search for individual EDCs using the chemical name or standard identifier. Physicochemical filter option in DEDuCT can be used to also filter EDCs based on their physicochemical properties such as molecular weight, number of hydrogen bond donors or acceptors, number of rotatable bonds. Chemical similarity filter gives the top 10 structurally similar EDCs in DEDuCT in comparison to the query molecule. **(B)** The Browse section in the web interface of DEDuCT can be used to view the EDCs based on the type of supporting evidence or their environmental source or chemical classification or systems-level perturbations and endocrine-mediated endpoints.

